# Heterosubtypic cross-reactivity of HA1 antibodies to influenza A, with emphasis on nonhuman subtypes (H5N1, H7N7, H9N2)

**DOI:** 10.1101/116327

**Authors:** Dennis E. te Beest, Erwin de Bruin, Sandra Imholz, Marion Koopmans, Michiel van Boven

## Abstract

Epidemics of influenza A vary greatly in size and age distribution of cases, and this variation is attributed to varying levels of pre-existing immunity. Recent studies have shown that antibody mediated immune responses are more cross-reactive than previously believed, and shape patterns of humoral immunity to influenza A viruses over long periods. Here we quantify antibody responses to the hemagglutinin subunit 1 (HA1) across a range of subtypes using protein microarray analysis of cross-sectional serological surveys carried out in the Netherlanc before and after the A/2009 (H1N1) pandemic. We find significant associations of responses, both within and between subtypes. Interestingly, substantial overall reactivity is observed to HA1 of avian H7N7 and H9N2 viruses. Seroprevalence of H7N7 correlates with antibody titer to A/1968 (H3N2), and is highest in persons born between 1954 and 1969. Seroprevalence of H9N2 is high across all ages, and correlates strongly with A/1957 (H2N2). This correlation is most pronounced in A/2009 (H1N1) infected persons born after 1968 who have never encountered A/1957 (H2N2)-like viruses. We conclude that heterosubtypic antibody crossreactivity, both between human subtypes and between human and nonhuman subtypes, is common in the human population.

## Introduction

Yearly epidemics of influenza A in temperate regions are notoriously variable in duration, size, and age distribution of cases [1–5]. This appears to be true for influenza pandemics caused by novel or reappearing subtypes as well [3, 5–10]. For instance, while mortality rates of 0.5%-1.0% have been reported for the 1918 pandemic, causing 20-100 million deaths worldwide, the recent 2009 pandemic was relatively mild, with an estimated overall attack rate of 24% and estimated mortality rate remaining below 0.01% [6, 9]. Varying levels of pre-existing immunity for the strain that is seeding the pandemic or yearly epidemic are driving the observed differences [1, 5, 7–9, 11–14]. Protection to influenza A infection and severity is mediated by cellular and humoral immune responses, with cellular responses being relatively conserved across viruses and in time, and antibody responses being more specific and variable [15, 16]. Recent studies, however, show that antibody-mediated immune responses, especially those directed against the stalk of the hemagglutinin protein, are more cross-reactive than previously believed [17–39].

To assess the extent of cross-reactivity of antibody responses to historic influenza viruses in the human population we analyze cross-sectional serological survey data from the Netherlands obtained before and after the A/H1N1 pandemic of 2009 [7]. The samples had been tested by hemagglutination inhibition test using pandemic A/2009 (H1N1) virus as antigen, and by hemagglutinin subunit 1 (HA1) protein microarray using a suite of human H1N1, H2N2, and H3N2 viruses, and avian H5N1, H7N7 and H9N2 viruses [40–42]. Earlier analyses focused on A/1918 (H1N1) and A/2009 (H1N1) HA1 antigens, to investigate diagnostic characteristics of the new tests to distinguish persons infected during the pandemic of 2009 from those with preexisting immunity [43], and to estimate infection attack rates and population levels of preexisting immunity [8]. The previous studies reveal that a strong correlation exists between the hemagglutination inhibition test and the HA1 microarray when using pandemic A/2009 (H1N1) virus as antigen, especially in samples of persons that were likely infected with the pandemic virus. Moreover, the microarray offers a sensitive and specific test to distinguish recently infected persons from those with pre-existing immunity. Here we take into account full response to 12 antigens in 357 persons, including those to HA1 antigens from avian H5N1, H7N7 and H9N2 viruses. We focus on correlations that must be caused by cross-reactive responses and cannot be caused by associations in infection histories, e.g., responses to avian influenza viruses, and responses against HA1 of A/1957 (H2N2) virus in persons born after 1968, i.e. who are too young to have been infected naturally.

## Materials and methods

The sera used in this paper originate from two age stratified serological surveys that were conducted before and after the influenza pandemic of 2009 [7]. The study was approved by the Medical Ethical Testing Committee of Utrecht University (Utrecht, the Netherlands), according to the Declaration of Helsinki (protocol 66-282/E). Written informed consent was given by participants (or next of kin/caregiver in the case of children) for suitably anonymized clinical records to be used in this study.

We draw an age stratified random subset of these surveys (167 pre-pandemic samples and 190 post-pandemic samples), oversampling sera with high A/2009 (H1N1) HI titers [43]. Persons who had received pandemic vaccinations were not eligible for selection because of the interference of vaccination with the microarray test results [41]. Full details on the antibody protein array have been described earlier [41, 42]. Briefly, antibody titers are calculated from the inflection point of twofold serial dilution series, with lower and upper dilutions of 1/20 and 1/2560, respectively. Throughout, a sample is said to be seropositive if it has a positive response in the first 1/20 dilution. Samples that are negative in the first 1/20 dilution are said to be seronegative. In the figures, these samples are assigned a titer value of 20, and in the statistical analyses these samples are considered to be left-censored.

For each person in the study, information on age, sex, and municipality of residence was available from the population register. The participants filled in a questionnaire, containing questions on demographic characteristics, living conditions, underlying illness, history of influenza-like illness, and seasonal influenza vaccination history in the preceding five years [7].

Samples are analyzed with nine representative antigens from human viruses and three avian isolates of main subtypes that are considered a risk of forward human transmission (H5N1, H7N7, H9N2). Specifically, we include five human H1N1 viruses (A/South Carolina/1/1918, A/WS/1933, A/New Caledonia/20/1999, A/Brisbane/59/2007, and the pandemic A/California/06/2009), one H2N2 virus (A/Canada/720/2005, a 1957-like virus), and three H3N2 viruses (A/Aichi/2/1968, A/Wyoming/2/2003, A/Brisbane/10/2007). Table 1 gives an overview of the antigens included.

**Table 1.** Antigens included in the microarray.

| Strain | Subtype |
| --- | --- |
| A/South Carolina/1/1918 | H1N1 |
| A/WS/1933 | H1N1 |
| A/New Caledonia/20/1999 | H1N1 |
| A/Brisbane/59/2007 | H1N1 |
| A/California/06/2009 | H1N1 |
| A/Canada/720/2005 (1957-like) | H2N2 |
| A/Aichi/2/1968 | H3N2 |
| A/Wyoming/2/2003 | H3N2 |
| A/Brisbane/10/2007 | H3N2 |
| A/Vietnam/1194/2004 | **H5N1** |
| A/Chicken/Netherlands/1/2003 | **H7N7** |
| A/Guinea fowl/Hong Kong/WF10/1999 | **H9N2** |

Overview of the origin of the HA1 antigens included in the antibody protein microarray (reproduced from [43]). Viruses of avian origin are denoted in bold.

Estimates of seroprevalence in the population are based on seropositivity of the test at the 1/20 or 1/40 dilution. In order to calculate population prevalence, weights are assigned to the individual sera to scale the data back to the population census and the population size of the HI strata. Confidence intervals for interval estimates of seroprevalence are calculated by the exact Clopper-Pearson binomial method. Smoothed antibody concentrations and smoothed seroprevalences are estimated with generalized additive models, using the gamlss procedure of the gamlss package. Pairwise associations of antibody responses are based on Kendall’s tau, a non-parametric correlation coefficient. For two measurements in two random subjects Kendall tau has a straightforward interpretation as the difference between the probability for the measurements to be in the same order and the probability of the measurements being in a different order. This implies that, for example, a value of tau=0.8 translates to a probability of (1-0.8)/2=0.1 that two measurements in two samples are in different order. Finally, linear regression analyses are performed using the statistical programming language R.

All data are available at the person-level on GitHub (github.com/mvboven/influenza-microarray-data). For each person we report sex, age in years, pre-versus post-pandemic sample, history of seasonal vaccination in the preceding five years, A/2009 (H1N1) HI titer, and microarray measurements for all 12 antigens. Also included are the population weights used to estimate population prevalence (see above).

## Results

### Group 1 viruses

Within group 1 influenza A viruses (H1, H2, H5, and H9), seroprevalence for the similar A/1918 (H1N1) and A/2009 (H1N1) hemagglutinins are high in both the pre- and postpandemic surveys in persons 20 years and older (Fig 1)[8, 43]. In persons under 20 years, seroprevalence is also high in the post-pandemic survey but not the pre-pandemic survey, an observation that has been exploited earlier to estimate infection attack rates [7, 8]. Seroprevalence for A/2007 (H1N1) is also high across all ages, and does not seem to differ between the pre- and post-pandemic surveys. Highest seroprevalence to A/2007 (H1N1) and highest antibody concentration measurements are observed in children who were 10-20 year old in 2009. These children likely had their first influenza infection(s) with A/2007 (H1N1)- like viruses. Seroprevalence and antibody titers to A/1957 (H2N2) virus (the Asian flu), which circulated from 1957-1968, are low in persons born after 1965 (who are unlikely to have been infected with H2N2), and much higher in persons born before 1965 [42]. Interestingly, high seropositivity (40%, 95%CI: 35%-45%) exists in the Dutch population against the avian H9N2 virus. Seropositivity of H9N2 is substantial across persons older than 10 years, and peaks in persons 40-60 years with estimated prevalence exceeding 50% when using a microarray titer cutoff of 20. Seropositivity of H9N2 is lower when using a higher microarray titer cutoff of 40, but remains high, especially in persons 40-60 years (>30%). Antibody responses to H5N1 virus are low overall (5.1%, 95%CI: 3.1%-7.9%), and are affected by history of seasonal influenza vaccinations (see below).

**Fig 1.**
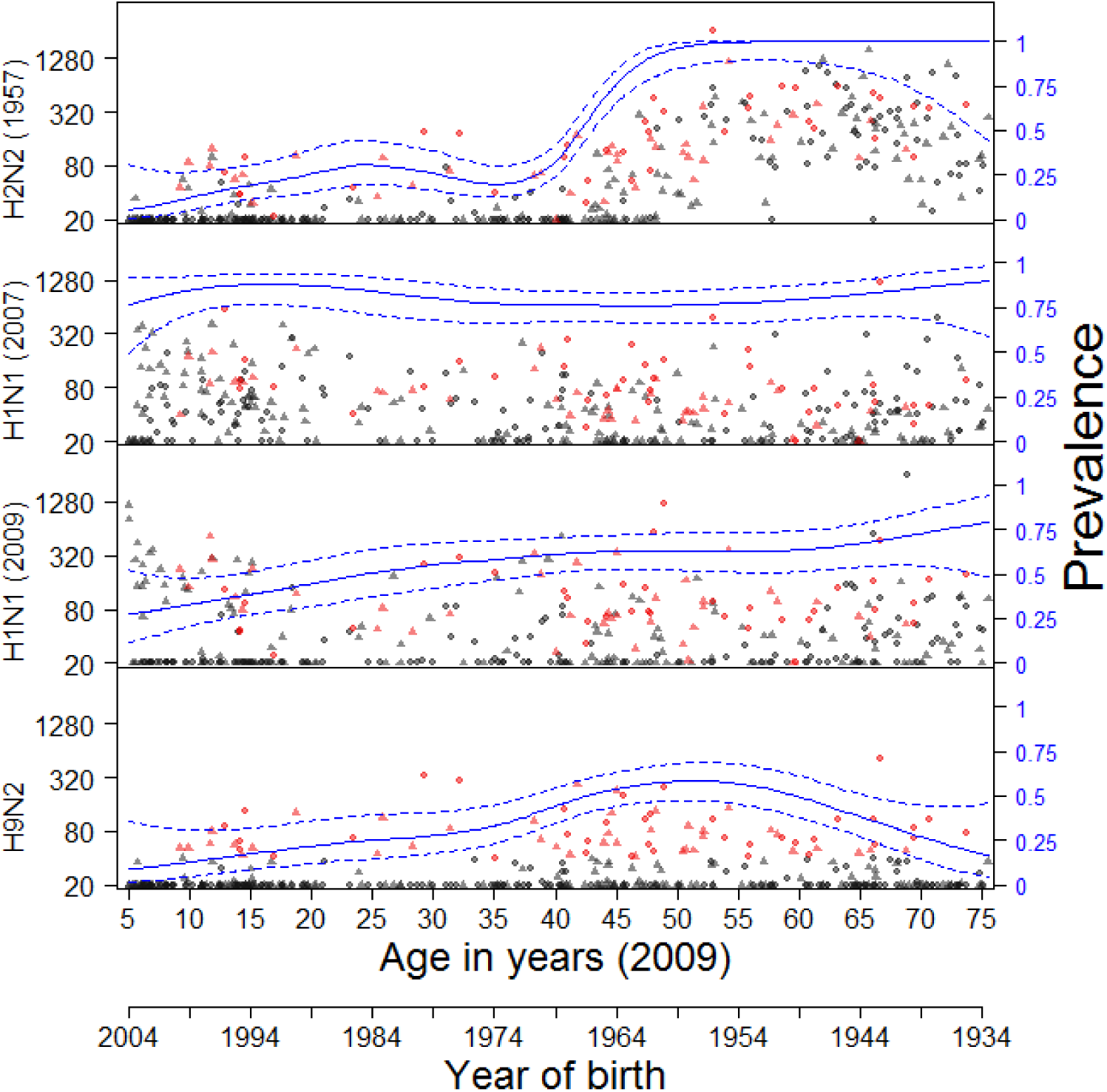
Age-specific antibody titers to group 1 influenza A viruses. Pre- and post-pandemic samples are represented by triangles and circles, respectively. Red dots indicate that H9N2 titer was ≥40. Solid and dotted lines show weighted seroprevalence estimates and associated 95% confidence ranges, estimated with a logistic generalized additive model (right axis). Seroprevalence estimates are weighted by their contribution to the population census [8, 43]. For clarity, only pooled pre- and post-pandemic prevalence estimates are presented. Antigens (HA1) included are from viruses A/South Carolina/1/1918 (H1N1), A/Canada/720/2005 (H2N2) (1957-like H2N2), A/Brisbane/59/2007 (H1N1), A/California/6/2009 (H1N1), A/Guinea fowl/Hong Kong/WF/10/1999 (H9N2).

Fig 2 shows antibody cross-correlations between 2009 pandemic H1N1 virus, the Asian flu (H2N2), and the avian H9N2 virus in different age groups. Antibody responses against H2N2 and H9N2 viruses are strongly correlated in both surveys, especially in persons under 45 years (tau=0.75 for both surveys, p<10−^4^ for both surveys). As these persons cannot have been infected naturally, these responses must result from a correlated response after infection with another virus. It is tempting to speculate that this is the pandemic H1N1 virus, as seropositivity against H2N2 or H9N2 in all but a handful of sera also implies seropositivity to the H1N1 virus. In young children (5-9 years) these patterns are even more pronounced. In this age group, all sera are seronegative (i.e. have no measurable antibodies) to the H1N1, H2N2, and H9N2 viruses in the pre-pandemic survey (n=15 samples), and 17 of 26 samples (65%) is seropositive in the post-pandemic survey. Of these, 6 are also positive for the H2N2 and H9N2 viruses. Similar results are observed in older children (10-19 years). In adults over 45 years who are likely to have experienced a natural H2N2 infection, the correlation between H2N2 and H9N2 is less pronounced, with low correlation coefficients of 0.18 and 0.24. In these persons, the H2 response is usually substantially higher than the H9 response.

**Fig 2.**
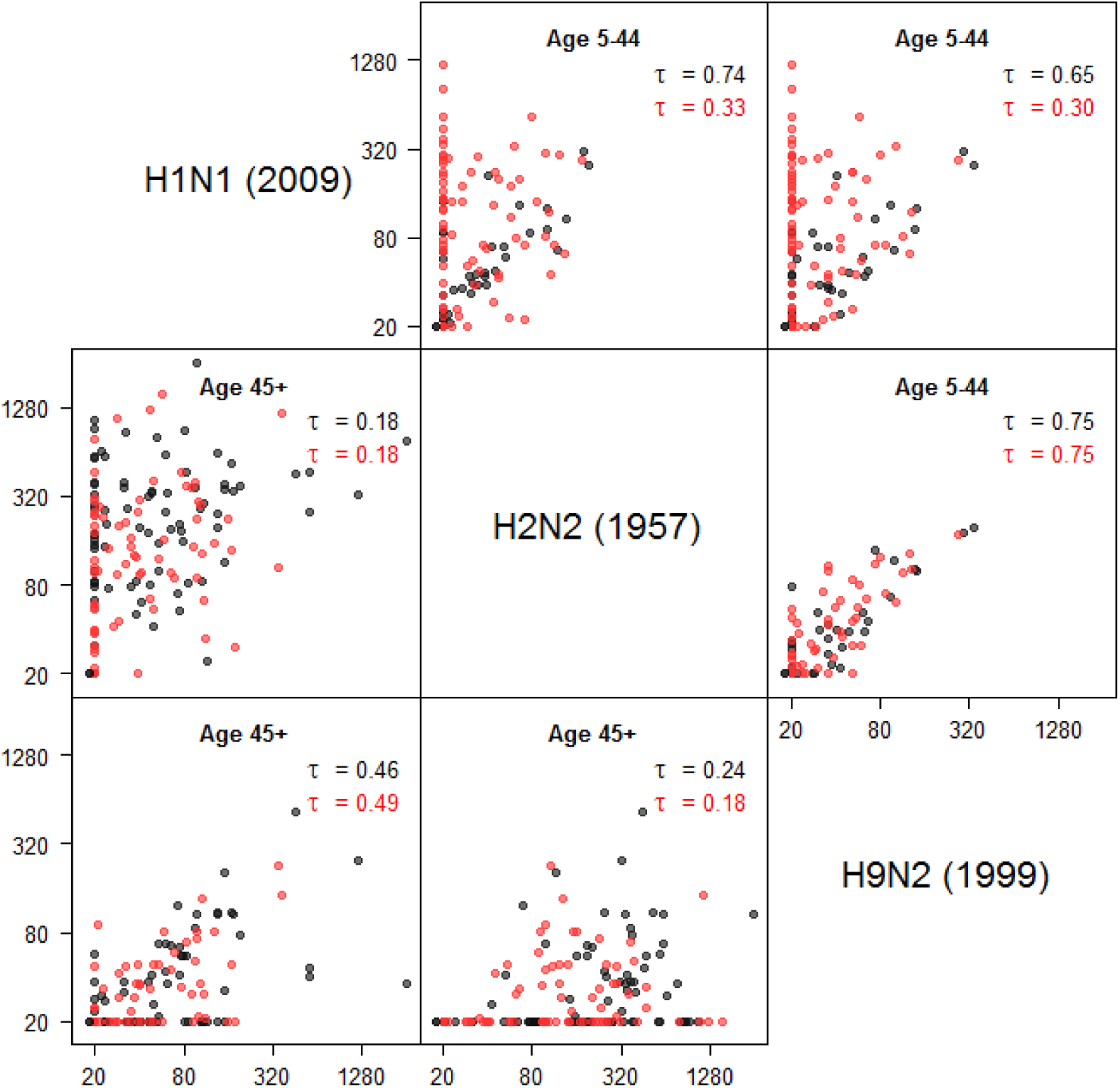
Correlations of antibody responses to group 1 influenza viruses. Results are given for age groups 5-44 years (above the diagonal) and 45+ years (below the diagonal). Black dots: prepandemic survey; red dots: post-pandemic survey. In each panel, Kendall’s tau is given for the pre- and post-pandemics surveys (black and red). Notice the strong positive correlation between H2N2 and H9N2 antibody titers in persons up to 45 years, i.e. persons who are unlikely to have experienced a natural H2N2 infection. Positive antibody titers to H2N2 and H9N2 are found almost exclusively in persons with positive A/2009 (H1N1) response, suggesting that A/2009 (H1N1) infection yields a response that is cross-reactive with both H2N2 and H9N2.

### Group 2 viruses

For group 2 influenza viruses (H3, H7), seroprevalence against H3N2 viruses is high across all age groups, reaching 100% in many age strata (Fig 3). Seronegative persons are only found in children (<20 years) with regard to their A/1968 (H3N2) response, and in adults over 40 years with regard to young H3N2 (2003, 2007) viruses. Antibody titers against A/1968 (H3N2) peak in persons born between 1954 and 1969. As children in the Netherlands contract their first influenza infection between 1 and 7 years [44], it is likely that a large fraction of persons born between 1954 and 1969 (and especially those born between 1961 and 1967) had their first influenza A infection(s) with 1968 H3N2-like viruses. The same holds for H3N2 (2003) virus, for which antibody titers are highest in persons born between 1989 and 1999. These persons likely had their first influenza infection with 2003 H3N2-like virus. Remarkably, seroprevalence against the avian H7N7 virus, albeit lower than against H9N2 virus, is substantial overall (population estimate: 10%, 95CI: 7.2%-14%%), and peaks in persons born between 1954 and 1969 (20%, 95%CI: 13%-28%). The peak coincides with the peak in A/1968 (H3N2) antibody titers.

**Fig 3.**
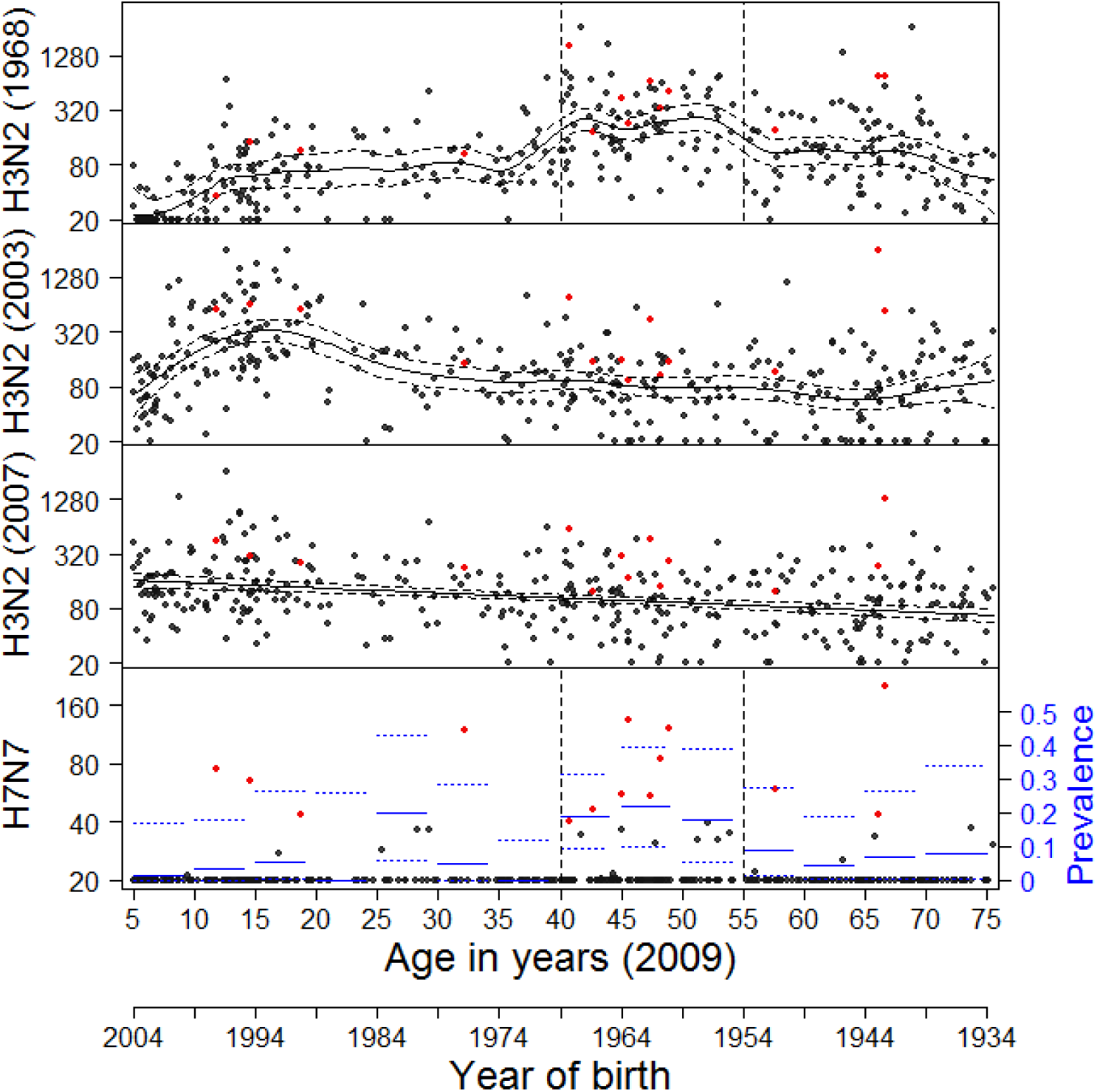
Age-specific antibody titers to group 2 influenza A viruses. Red dots indicate an H7N7 titer ≥40. Seroprevalence to H3N2 viruses reaches 100% in most age strata. Solid and dotted lines show the mean weighted antibody concentrations and associated 95% confidence ranges, estimated with a generalized additive model. For H7N7, solid and dotted lines show interval population-weighted seroprevalence estimates and associated 95% confidence bounds (right axis). Antigens (HA1) included are from viruses A/Aichi/2/1968 (H3N2), A/Wyoming/2003 (H3N2), A/Brisbane/10/2007 (H3N2), and A/Ck/Netherlands/2003 (H7N7).

### Correlations between avian viruses

Fig 4 shows correlations of antibody responses to avian viruses (H5N1, H7N7, H9N2). Interestingly, there is a strong correlation between the H9N2 and H7N7 responses, especially in older persons, even though these viruses belong to different influenza A subtype groups (group 1 versus group 2). Seropositivity to H5N1 is low overall, as only a handful of samples (n=21) have measurable antibody titers to this virus. There is no noticeable correlation between H5N1 and H9N2, or between H5N1 and H9N2. Analysis of the data by vaccination status in the preceding five years shows that seasonal influenza vaccination has an impact on H5N1 antibody titers. In fact, a linear regression model explaining H5N1 antibody titers by age and vaccination status (0 vaccinations versus **>**≥1 seasonal influenza vaccinations in the preceding five years) shows that vaccinated persons have significantly higher H5N1 antibody titer than unvaccinated persons (p=0.0095).

**Fig 4.**
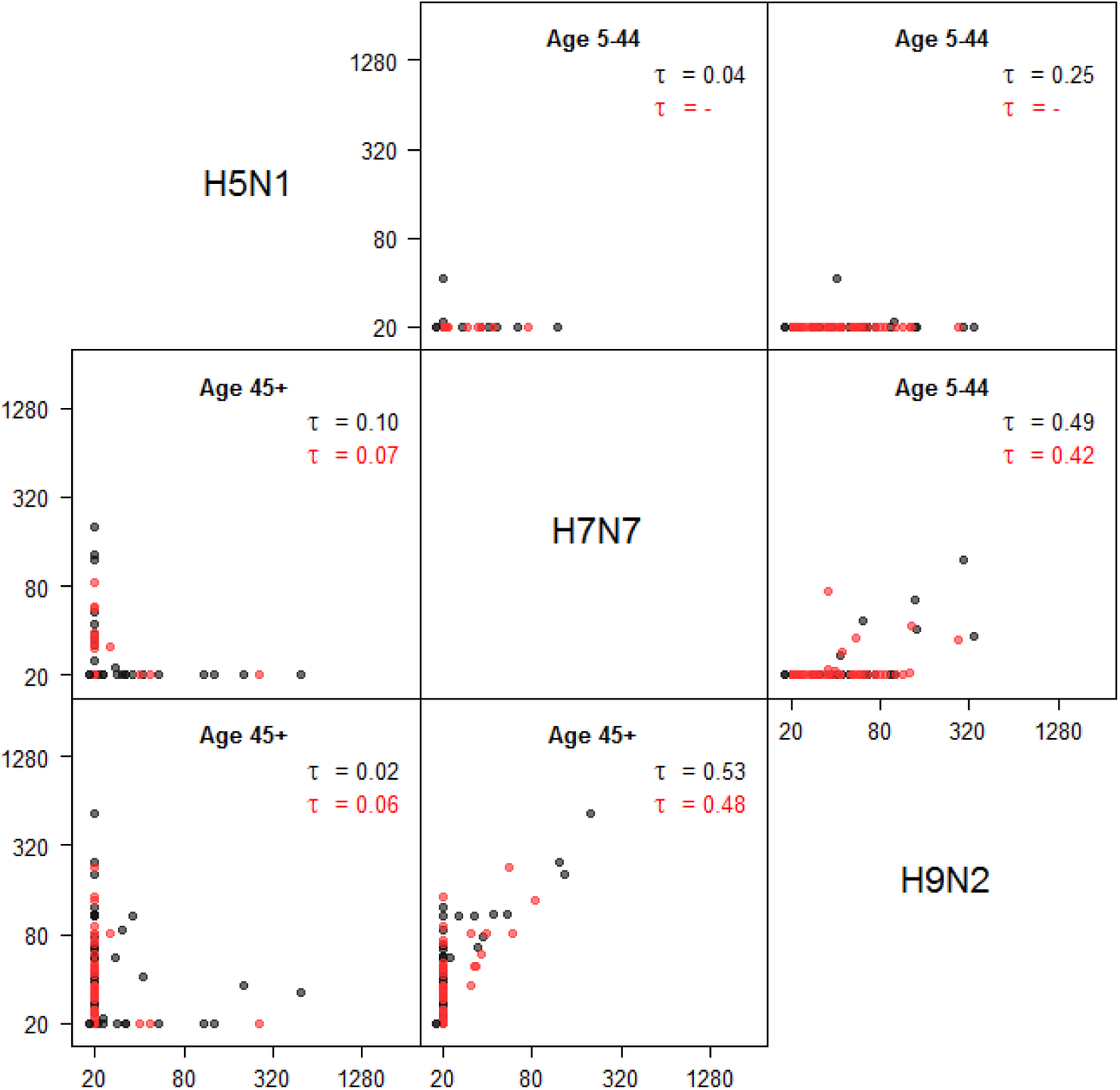
Correlations of antibody responses to avian influenza A viruses. Results are given for age groups 5-44 years (above the diagonal) and 45+ years (below the diagonal). Black dots: pre-pandemic survey; red dots: post-pandemic survey. In each panel, Kendall’s tau is given for the pre- and post-pandemics surveys (black and red). Notice that all persons with positive H7N7 response also have a positive H9N2 response. Antigens included are from viruses A/Vietnam/1194/2004 (H5N1), A/Chicken/Netherlands/1/2003 (H7N7), and A/Guinea fowl/Hong Kong/WF 10/1999 (H9N2).

## Discussion

Correlations in the antibody responses to different influenza viruses may arise by crossreactivity of responses after infection by one virus, or by associations in infection histories (e.g., it is more likely that you are infected with virus Y if you have been infected with virus X). In general, it is difficult to distinguish between the two explanations. In some cases, however, antibody response associations can only be the result of cross-reactivity, either because the virus of interest has not circulated in the human population, or because the virus has not been in circulation in certain age strata (H2N2 in persons born after 1968, A/2009 (H1N1) in the pre-pandemic serological survey). Here we have exploited these observations to explore cross-reactivity of antibody responses to influenza A viruses in a (structured) random sample from the Dutch population.

Our analyses have uncovered 1) that HA1 antibodies that react to avian H7N7 and H9N2 viruses are present in a significant part of the Dutch population, 2) that infection with 2009 pandemic H1N1 virus often induces measurable antibodies to HA1 of H2 and H9 viruses, and 3) that seroprevalence to H7 virus is high in persons born between 1954 and 1969. Our results are in line with earlier finding of ‘original antigenic sin’ and ‘antigenic seniority’, i.e. antibody responses are generally strongest to those viruses encountered early in life, and with the related recent concept of ‘antigenic landscapes’, i.e. antibody responses after primary infection extending beyond the extent of intrinsic cross-reactivity [1, 11–13, 30, 38, 39, 45].

All results reported here are based on a protein microarray test based on the HA1 subunit of the hemagglutinin. As with the hemagglutination inhibition assay, measurements based on this test are not necessarily protective. It is known, however, that responses in the microarray correlate well with corresponding HI titers [41–43]. In our data, we found a strong correlation between microarray titers and (standardized) HI titers when using A/2009 (H1N1) antigen, with a (standardized) HI titer of 40 translating roughly to a microarray titer of 80 [43]. It is also known that there is a positive relation between HI titers and protective immunity, and it is sometimes stated that a HI titer of 40 would translate to a 50% probability of protective immunity [46]. If the above reasoning would prove true, one could argue that A/2009 (H1N1) microarray titer of 80 or more is expected to be protective in 50% of persons. Supporting this line of reasoning is provided by the observation that the HA1 subunit includes the immunodominant globular head of the hemagglutinin, which contains the four (or five) main antibody epitopes that are traditionally believed to provide sterilizing immunity.

We included three avian influenza viruses in this study that belong to nonhuman subtypes (H5N1, H7N7, H9N2). These viruses have caused numerous human infections, and might develop into fully human transmissible viruses. It may seem plausible that there will be little antibody-mediated pre-existing immunity in the population to these viruses (but see e.g., [33, 47]). If one would extend the above lines of reasoning for A/2009 (H1N1) to other antigens, one could argue that there would indeed be limited immunity to H7 viruses in the Dutch population, as only a handful of participants have a microarray titer ≥80, mainly in the persons born between 1954 and 1969 (Fig 3). For H9, on the other hand, a substantial fraction (30/357-8.4%) of the population has high microarray titers of >≥80, which could translate into protective immunity. For H5, seroprevalence is very low (5.1% at a microarray titer threshold of 20), and modulated by seasonal vaccination. Interestingly, however, the correlation of H5N1 in older persons (over 44 years) is strongest with the A/1918 (H1N1) and the A/1933 (H1N1) antigens, with Kendall correlations of tau=0.33 and tau=0.46, respectively. Together, these finding support the recent observation that the incidence of severe H5N1 and H7N9 infections is lower than expected in age groups that had their first infections with viruses from the same antigenic family (i.e. H1/H2 in case of H5N1 and H3 in case of H7N9) [48].

Finally, our results shed some light on recent studies on broadly reactive and broadly neutralizing antibodies on the hemagglutinin protein, in particular on HA2 but to a lesser extent also on HA1 [17–32, 34]. These studies have mostly used animal models and laboratory experiments to study heterosubtypic antibody responses. Here we have added that these findings are not only found in animals in the laboratory and selected populations (e.g., [34, 47]), but that cross-reactivity can be estimated quantitatively in the human population. Our data thus lend additional evidence to the hypothesis that as humans age and accumulate infections, antibody responses become increasingly directed to more conserved parts of the hemagglutinin, and thereby more cross-reactive [11, 12, 18, 30].

## Acknowledgments

This work was supported by the National Institute for Public Health and the Environment project S/210096. The funders had no role in study design, data collection and analysis, decision to publish, or preparation of the manuscript.

## Author contributions

DtB, MK, and MvB designed the study. Experimental analyses were performed by EdB and SI. DtB performed the analyses. DtB and MvB wrote the draft manuscript text. All authors reviewed the manuscript.

